# Cardiovascular risk gene *HDAC9* drives maladaptive vascular remodeling after arterial injury

**DOI:** 10.64898/2026.05.12.723753

**Authors:** Federica Tosato, Donovan Correa-Gallegos, Arailym Aronova, Remco T. A. Megens, Christian Behrends, Yaw Asare

## Abstract

Arterial restenosis following balloon angioplasty – a procedure performed to re-establish vessel patency in atherosclerotic cardiovascular disease – remains a major clinical challenge and a key barrier to durable revascularization. Endothelial denudation induced by angioplasty triggers an inflammatory cascade that drives vascular smooth muscle cell (VSMC) proliferation, migration, and phenotypic switching, culminating in neointimal hyperplasia and restenosis^1^. Human genetics-guided target discovery has proven more effective than non-guided approaches in revealing causal pathways of complex cardiovascular traits^2^. Genetic variants at *Histone Deacetylase 9* (*HDAC9*) are a major risk factor for cardiovascular disease^3,4^ and is associated with increased carotid intima–media thickness and modulation of VSMC phenotype^4^. Here, using an experimental model of arterial injury that faithfully mirrors the vascular response to balloon angioplasty in humans, we show that HDAC9 drives maladaptive remodeling of the arterial wall following vascular injury.

## Results

All data are presented in Figure 1. Sequencing and proteomics datasets have been deposited in the repositories GSE312075 and PXD071212, respectively. To assess the role of HDAC9 in arterial restenosis, *Hdac9*^*–/–*^*Apoe*^*–/–*^ and *Hdac9*^*+/+*^*Apoe*^*–/–*^ mice were subjected to wire-induced injury of the common carotid artery. Hypercholesterolemia, a common comorbidity in atherosclerotic patients, was induced by feeding an atherogenic diet one week before endothelial denudation. Neointima formation was evaluated 3 weeks after injury. Histomorphometric analysis of movat pentachrome-stained sections revealed a significant reduction in neointimal area in *Hdac9*-deficient mice relative to controls (**Fig. 1Ai, Aii**). Blood leukocyte counts, total cholesterol, and body weight were comparable between genotypes, indicating that the attenuated neointima formation was mediated by vascular rather than systemic factors. Quantification of the neointimal cellular composition showed a reduced cell number, attributable to fewer VSMCs in *Hdac9*^*–/–*^*Apoe*^*–/–*^ (**Fig. 1Aiii, Aiv**). Given the pivotal role of endothelial cell (EC) activation in initiating the inflammatory cascade that drives VSMC proliferation and migration during neointima formation, we examined the impact of Hdac9 on endothelial activation in arteries from hyperlipidemic mice. Using two-photon imaging, we quantified Vcam-1 expression – a marker of EC activation – in freshly isolated and *ex vivo* mounted carotid arteries (80 mm Hg) from both genotypes after four weeks of atherogenic diet. *Hdac9*-deficient carotid arteries exhibited markedly reduced Vcam-1 expression compared with controls (**Fig. 1Bi, Bii**), indicating reduced endothelial activation. To explore the underlying molecular pathways, we performed quantitative proteomic profiling of Hdac9-depleted mouse aortic endothelial cells (MAoECs) stimulated with TNF-α, an endogenous driver of sterile vascular inflammation. Hdac9 loss perturbed multiple inflammatory pathways, including MAP kinase activation, TRAF6-mediated induction of NF-κB and MAP kinases, and activation of AP-1 transcription (**Fig. C**) consistent with the diminished endothelial activation observed in *Hdac9*^−/–^ carotid arteries.

**Fig. 1.**
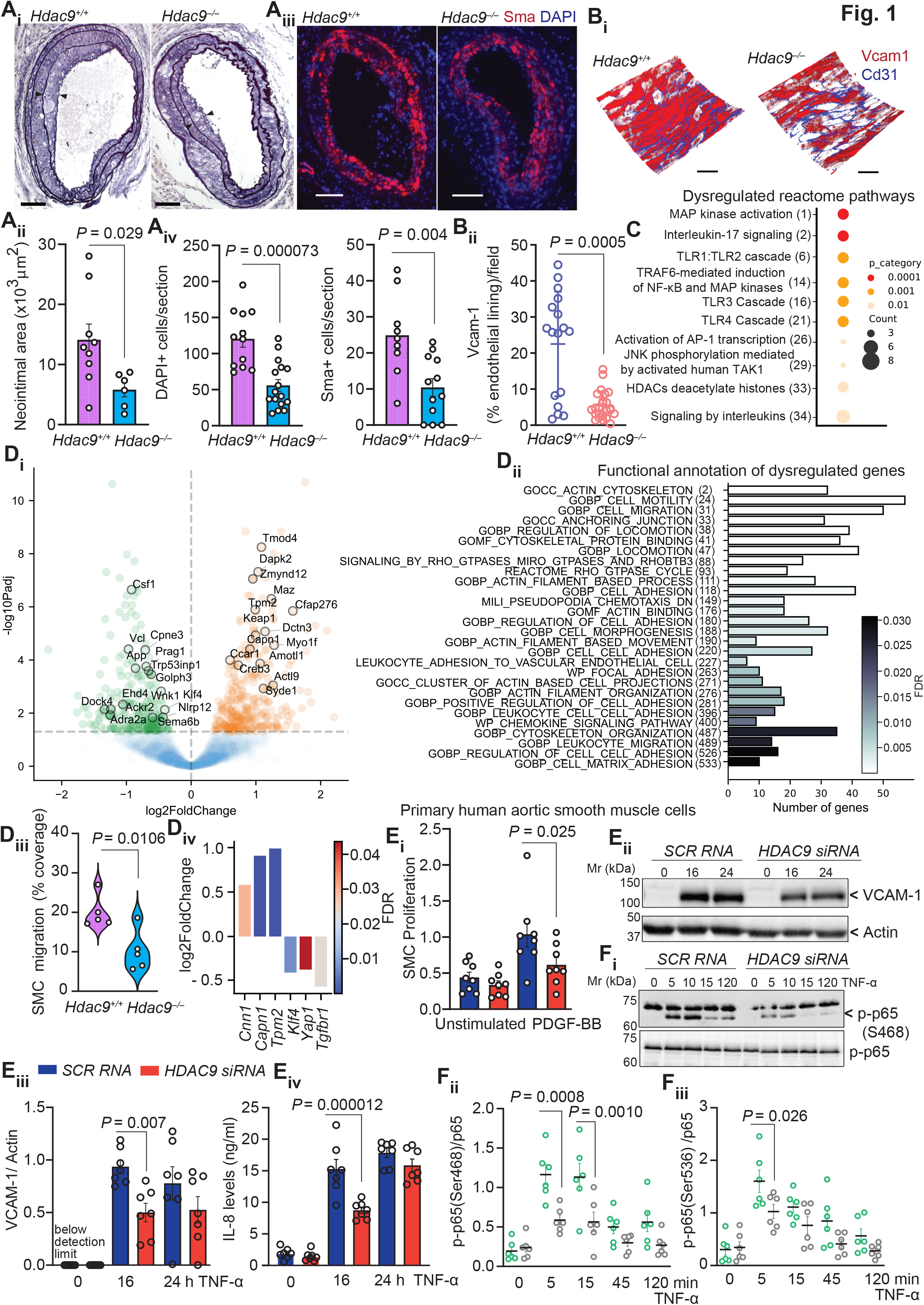
HDAC9 drives maladaptive vascular remodeling after endothelial denudation. **Ai-Aiv**, Wire injury model. *Hdac9*^*–/–*^*Apoe*^*–/–*^ and *Hdac9*^*+/+*^*Apoe*^*–/–*^ control littermate mice received Western-type diet one week before and for another three weeks after injury. **Ai**, Neointima formation was assessed in movat pentachrome-stained sections three weeks after injury. Scale bar represents 100 µm. **Aii**, Quantification of neointima area. n = 6-9 mice per genotype. Two-sided unpaired t test. **Aiii**, Representative DAPI and α-Sma staining. **Aiv**, Quantification of total cells in the neointima, and VSMC content. n = 3-4 mice per genotype. The quantification of cells is presented as the percentage of positively stained cells per section. Two-sided unpaired t test and Two-sided Mann-Whitney test. Scale bar represents 100 µm. **Bi**, Two-photon microscopy of endothelial Vcam-1 expression in *ex vivo* mounted carotid arteries. *Hdac9*^*–/–*^*Apoe*^*–/–*^ and *Hdac9*^*+/+*^*Apoe*^*–/–*^ control littermate mice received Western-type diet for 4 weeks followed by imaging of Vcam-1 and Cd31 (n = 6-7 mice per group). **Bi**, Quantification of endothelial Vcam-1 in mounted carotid arteries. Scale bar, 50 µm. The quantification of Vcam-1 expression is presented as the percentage of endothelial lining per field of view. **C**, Quantitative proteomic profiling of MAoEC. Shown is the reactome pathway of differentially regulated proteins in TNF-α-stimulated cells (FDR-adj. p value < 0.05). **Di, Dii**, Transcriptomic profiling of mouse aortic smooth muscle cells (MAoSMC). MAoSMC, isolated from *Hdac9*^*–/–*^ *Apoe*^*–/–*^ and *Hdac9*^*+/+*^*Apoe*^*–/–*^ control mice, were stimulated with TNF-α for 6 hours and subjected to gene expression profiling. **Di**, Volcano plot of log_2_FoldChange (*Hdac9*^*+/+*^*Apoe*^*–/–*^ versus *Hdac9*^*–/–*^*Apoe*^*–/–*^) and –log_10_p-adjusted values showing top 30 differentially expressed genes related to cell migration. Highlighted in green and orange are the upregulated and downregulated genes respectively. **Dii**, Gene set enrichment analysis showing functional annotation of dysregulated genes (all FDR corrected). **Diii**, Scratch wound-healing assay. VSMC, isolated from *Hdac9*^*–/–*^*Apoe*^*–/–*^ and *Hdac9*^*+/+*^*Apoe*^*–/–*^ control mice, were stimulated with PDGF-BB for 24 hours and analyzed for their migratory capacity. Quantification of the area covered by migrated cells. n = 5 mice per genotype. Two-sided unpaired t test. **Div**, Contractile markers and modulators of VSMC phenotypic transformation from the transcriptome analysis. **Ei-Fiii**, Functional and molecular assays in human aortic smooth muscle cells (HAoSMC). HAoSMC were transiently transfected with *HDAC9* siRNA or scrambled control (*SCR*) RNA for 72 hours and subsequently stimulated with TNF-α (20 ng/mL) for indicated periods or left untreated. **Ei**, Quantification of PDGF-BB-induced proliferation of HAoSMCs by EdU assay. **Eii**, Representative immunoblots of VCAM-1 and Actin. Quantification of VCAM-1 normalized to Actin levels (**Eiii)**, and IL-8 release (**Eiv). Fi**, Representative immunoblots of p65 phosphorylation at serine 468 and total p65 levels. Quantification of phosphorylated p65 at serine 468 (**Fii**) and serine 536 (**Fiii**) normalized to total p65. n = 6-7 independent experiments. Two-way ANOVA with Bonferroni multiple comparison test for comparison of *HDAC9* siRNA vs. *SCR* RNA. Data are means ± SEM.

Endothelial damage and activation results in key signals that regulate VSMC function critical in arterial wall remodeling. Hence, we analyzed the transcriptome of mouse aortic smooth muscle cells (MAoSMC) isolated from *Hdac9*^*–/–*^*Apoe*^*–/–*^ and control *Hdac9*^*+/+*^*Apoe*^*–/–*^ mice. After checking the purity, cells were stimulated with TNF-α and subjected to bulk RNA sequencing. Gene set enrichment analysis revealed differential expression of genes associated with cell adhesion, cell motility, and migration, in *Hdac9*-deficient cells (**Fig. 1Di, Dii**). Consistent with these findings, scratch wound assays demonstrated markedly reduced migratory capacity of *Hdac9*-deficient VSMCs compared with controls (**Fig. 1Diii**). Transcriptomic data further indicated a retention of a contractile phenotype, characterized by upregulation of the contractile markers Cnn1, Capn1, and Tpm2, and downregulation of key modulators of VSMC phenotypic transformation, including Klf4, and Yap1 (**Fig. 1Div**). In primary human aortic smooth muscle cells (HAoSMC), siRNA-mediated knockdown of *HDAC9* diminished HAoSMC proliferation (**Fig. 1Ei**). Mechanistically, *HDAC9* depletion attenuated TNF-α-induced pro-inflammatory responses in HAoSMC, evidenced by reduced VCAM-1 expression, as well as decreased IL-8 production (**Fig. 1Eii-Eiv**). These results mirror the improved arterial remodeling observed in *Hdac9*-deficient carotid arteries (**Fig. 1Ai, Aii**,). Because the transcriptional changes in both *Hdac9*-deficient arteries and *HDAC9*-silenced HAoSMCs resembled NF-κB-driven signatures, we next investigated the role of HDAC9 in NF-κB signaling. TNF-α–induced p65 phosphorylation was markedly reduced in *HDAC9*-silenced HAoSMCs, predominantly at serine 468 and, to a lesser extent, serine 536 (**Fig. 1Fi-Fiii**). This phosphorylation pattern contrasts with our previous findings in macrophages and endothelial cells, where HDAC9 loss primarily affected phosphorylation at serine 536. Because site-specific phosphorylation of p65 encodes gene-selective transcriptional outputs, these results - together with our earlier work^5^ - support the NF-κB ‘barcode’ concept and reveal a cell type-specific activating role of HDAC9 in NF-κB signaling that requires further detailed investigation beyond the scope of the current study.

Collectively, we show that (i) genetic deletion of *Hdac9* attenuates neointima formation upon arterial injury in hyperlipidemic mice; ii) *Hdac9* deficiency mitigates endothelial and smooth muscle cell inflammatory responses, promoting a contractile phenotype and preventing a migratory state in VSMCs; and iii) HDAC9 may function as a molecular switch that modulates NF-κB activation in a cell type-specific manner. Our findings identify HDAC9 as a potential therapeutic target in pathological arterial remodeling following percutaneous coronary intervention.

## Acknowledgements

We are grateful to Dr. Yury Bokov for the help with the HAoSMC experiments and to Dr. Laura Strohm for her expertise on quantitative proteomics. We thank Dr. Aditi Methi for the help with data management and Melanie Schneider for her help with histology. This work was supported by grants from the Deutsche Forschungsgemeinschaft (DFG; CRC 1123 [B3], AS 575/1-1, AS 575/2-1) and Fritz Thyssen Foundation (10.24.2.001MN) to Y.A. R.M. received support from DFG (SFB1123-Z1, INST409/97-1 FUGG).

## Contributions

Y.A. conceived and designed research, with input from F.T; F.T., A.A., R.M. and Y.A., performed research; F.T., D.C-G., R.M., C.B, and Y.A., analyzed data; Y.A. and F.T. wrote the paper with input from all authors.

## Ethics declarations

Competing interests

The authors declare no competing interests.

Ethics

The Institutional Animal Care Committee of Regierung von Oberbayern (ROB-55.2-2532.Vet_02-14-187) approved all animal experiments.

## Supplementary Material

### Materials and Methods

#### Mouse model of arterial wire-induced injury

*Hdac9*^*–/–*^*Apoe*^*–/–*^ mice were generated as previously described^1^. Mice had ad libitum access to food and water and were housed in a specific pathogen-free animal facility under a 12 h light-dark cycle. 6-8 weeks old female *Hdac9*^*–/–*^*Apoe*^*–/–*^ and control littermate *Hdac9*^*+/+*^*Apoe*^*–/–*^ mice, placed on an atherogenic western-type diet (21% fat, 0.15% cholesterol) for 1 week, were anesthetized with Medetomidine (0,5 mg/kg), Midazolam (5 mg/kg) and Fentanyl (0,05 mg/kg) and subjected to wire-induced arterial injury of the common carotid artery as established^2^. Briefly, after midline neck incision, the left external carotid artery was tied off distally, and via transverse arteriotomy, a 0.014 inch flexible angioplasty guide wire was advanced by 1 cm. Complete and uniform endothelial denudation was achieved by three passes along the common carotid artery with a rotating motion. Thereafter, mice were placed on western-type diet for another 3 weeks and neointima formation was assessed. Animals with broken common carotid artery sections were excluded before immunohistochemistry. Cholesterol and triglyceride levels in blood serum were quantified using enzymatic assays (Cayman) according to the manufacturer’s protocol. All animal experiments and data analysis were performed under blind conditions for the genotype and were approved by the local ethics committee.

#### Two-photon imaging of whole-mount tissue

6-8 weeks old *Hdac9*^*–/–*^*Apoe*^*–/–*^ and control littermate *Hdac9*^*+/+*^*Apoe*^*–/–*^ mice received western-type diet for 4 weeks and carotid arteries were explanted and mounted on glass micropipettes^3-5^. CD31 and VCAM-1 expression were quantified using CD31 (#48031182, eBioscience) and VCAM-1 (#105724, Biolegend) antibodies. The samples were imaged using a LeicaSP5IIMP upright two-photon laser scanning microscope with a Ti:Sapphire Laser (Spetra Physics MaiTai Deepsee HP) tuned at 800nm and a 20×NA1.00 (Leica) water dipping objective. Image acquisition and processing was performed using LasX software (Leica).

#### Immunohistochemistry

For analyzing neointima formation after wire-induced arterial injury, 5 µm thick section of PFA-fixed, paraffin-embedded carotid arteries were stained with Movat’s pentachrome. Neointimal areas were quantified in serial sections (5 μm; 10-12 per mouse) within 500 μm from the bifurcation and planimetry of the areas within external elastic lamina, internal elastic lamina, or lumen using Image J analysis software. Smooth muscle cells (SMCs) in neointimal plaques were visualized by immunofluorescent staining for Sma-cy3. Nuclei were counterstained by 4’, 6-diamidino-2-phenylindol (DAPI). All images were recorded with a Leica DMLB fluorescence microscope and CCD camera, and quantification of lesion size and composition was performed using Image J analysis software. All analyses were performed without prior knowledge of the genotype.

#### Isolation of mouse aortic smooth muscle cells

Mouse aortic smooth muscle cells (MAoSMCs) were freshly isolated from entire mouse aorta. After removing the adventitia, the aorta was dissociated in 3 mg/ml of Collagenase Type II (#C2-22-1G, Merck) dissolved in 1.5 mL DMEM 1X + GlutaMAX (#31966-021, Gibco) containing 10 % FBS and 50 mg/mL gentamycin, and placed for one hour at 37°C in shaking. Afterwards, the cells were centrifuged for 5 minutes at 300 g and the pellet resuspended in full DMEM medium and seeded on collagen G-coated (#L7213, Sigma-Aldrich) wells. The purity was determined by smooth muscle cell actin (#C6198, Sigma-Aldrich) and Transgelin (#ab10135, Abcam) expression by immunocytochemistry and imaging was performed with the confocal microscope (LSM 880, Zeiss) using the 10× objective and analyzed with the ZEN software (Zeiss).

#### Bulk RNA sequencing

MAoSMCs were isolated from *Hdac9*^*-/-*^*Apoe*^*-/-*^ *and Hdac9*^*+/+*^*Apoe*^*–/–*^ mice. At passage 3, the cells were seeded in 6 cm dish and stimulated for 6 hours with TNF-α (20 ng/mL, #315-01A, Peprotech) or left untreated. Following one hour serum starvation, cells were lysed with RLT buffer containing β-mercaptoethanol and total RNA isolated with the Maxwell RSC48 instrument and miRNA Tissue Kit (Promega, Fitchburg, USA). The Agilent Bioanalyzer (Agilent RNA 6000 Nano kit, Agilent Technologies, Santa Clara, CA, USA) was used to assess the RNA quality. Libraries were then prepared using the Lexogen CORALL mRNA-seq Library Prep Kit V2 (Lexogen, Vienna, Austria) at half the volume recommended in the manual. Briefly, 200 ng of total RNA were captured on oligo-dT beads and reverse transcribed directly on the beads, followed by the synthesis of the second cDNA strand after adapter ligation. The adapted cDNA was eluted and then amplified with barcoded primer. The finalized, barcoded libraries were pooled and sequenced on an Illumina NextSeq 2000 (Illumina, San Diego, CA, USA) with a read length of 60 nucleotides in paired-end mode. For sequence analysis, raw paired-end RNA-seq reads were quality-checked using FastQC (v0.12.0) and aligned to the Mus musculus reference genome (GRCm38) using HISAT2 (v2.2.1) with default parameters. Gene-level quantification was performed with featureCounts (Subread v2.0.2), assigning uniquely mapped reads to annotated exons based on Ensembl gene models. The resulting count matrix was imported into Python, where lowly expressed genes (<100 total counts across all samples), ribosomal, mitochondrial, hemoglobin, and non-annotated gene entries (e.g., ENSMUS…, Rik genes) were removed. Differential expression analysis was conducted using PyDESeq2 (v0.5.0), including normalization and model fitting. Differentially expressed genes were defined using Wald tests with Benjamini–Hochberg correction, applying an adjusted p-value threshold of <0.05. Functional annotation of dysregulated genes was generated by GSEA. Transcriptomic data are available online under the accession number: GSE312075.

#### Quantitative proteomics

Mouse aortic endothelial cells (MAoECs) were transfected with predesigned ON-TARGETplus SMARTpool human *HDAC9* siRNA or non-targeting control by Lipofectamine RNAiMax Transfection Reagent (Invitrogen). Following 72 hours of incubation, cells were stimulated with 20 ng/ml human TNF-α or left untreated. Thereafter, cells were collected and lysed in urea buffer (9 M Urea, 50 mM Tris pH 8, 150 mM NaCl, 1x Roche protease inhibitor cocktail) followed by short sonification and samples processed as previously described^6^. MS raw data were processed with MaxQuant (version 1.6.0.1). The mass spectrometry proteomics data have been deposited to the ProteomeXchange Consortium via the PRIDE^7^ partner repository with the dataset identifier PXD071212.

#### Primary cell culture, transfection and gene silencing

Human Aortic Smooth Muscle Cells (HAoSMCs) were purchased from PromoCell (Heidelberg, Germany), plated on cell culture dishes coated with collagen (Biochrom AG, Berlin, Germany) and cultured in smooth muscle cell growth medium (PromoCell, Heidelberg, Germany), according to manufacturer’s recommendations. Transfection of HAoSMCs with predesigned ON-TARGETplus SMARTpool human *HDAC9* siRNA or non-targeting control (Dharmacon) was performed by electroporation using the HAoSMCs Nucleofector Kit (Lonza, Cologne, Germany) following the manufacturer’s instructions. Cells were then allowed to recuperate for 72 hours before being used in the experiment. HAoSMCs were stimulated with 20 ng/ml human TNF-α (PeproTech, Rocky Hill, NJ, USA) for the indicated time periods.

#### Scratch Wound Healing Assay

MAoSMCs, isolated from *Hdac9*^*-/-*^*Apoe*^*-/-*^ *and Hdac9*^*+/+*^*Apoe*^*–/–*^ were seeded in Collagen G-coated 24-well plates and allowed to form a confluent monolayer. Subsequently, a wound was generated using a 1000-μL pipette tip, and the cells were stimulated with PDGF-BB (10 ng/mL, #315-18, Peprotech). Images of the same area were taken at 0 and 24 hours, and the percentage of coverage was determined using ImageJ software.

#### Cell lysis and western blot analysis

For total cell lysates, HAoSMCs were washed with cold PBS and lysed with 1x NuPAGE-LDS-sample buffer (Invitrogen) containing 1 mmol/l DTT (Sigma Aldrich, Hamburg, Germany). Total cell lysates were separated by SDS-PAGE, transferred to a PVDF membrane (Whatman GmbH), and detected with the appropriate antibodies. Primary antibodies were incubated overnight at 4°C. As secondary antibodies, HRP-conjugated anti-mouse or anti-rabbit antibodies were used, and blots were developed with Immobilon Western HRP Substrate (Merck Millipore). Protein bands were visualized with a Fusion Fx7 and quantified using Image J 1.47v software (Wayne Rasband).

#### ELISA

HAoSMCs were transfected with predesigned ON-TARGETplus SMARTpool human *HDAC9* siRNA or non-targeting control (Dharmacon) as above. After 72 hours of incubation, cells were stimulated with 20 ng/ml human TNF-α for the indicated period. The secretion of chemokine IL-8 by HAoSMCs were quantified using commercially available ELISA kits (Invitrogen).

#### Flow cytometry

EDTA-buffered blood, obtained by retro-orbital puncture, was treated with erythrocyte lysis buffer (0.155 M NH_4_Cl, 10 mM NaHCO_3_). All cell suspensions were carefully washed and stained with FACS staining buffer and combinations of antibodies to Cd45, Cd11b, Cd3, Cd4, Cd8a, Cd25, Cd11c, B220, Ly6G (eBioscience), and Ly6C (Miltenyi Biotec). Flow cytometric analysis was performed using FACSVerse and FACSuite software (BD Biosciences) after appropriate fluorescence compensation, and leukocyte subsets were gated using FlowJo software (Treestar). B-cells were identified as Cd45^+^B220^+^; T-cells as Cd45^+^Cd3^+^; neutrophils as Cd45^+^CD11b^+^Ly6G^+^; monocytes as Cd45^+^Cd11b^+^Ly6C^+^.

#### Statistical analysis

Statistical analyses were performed with Graphpad Prism 10 using two-tailed unpaired T-test, Mann-Whitney test, or two-way Anova with Bonferroni’s multiple comparison’s test as appropriate after normality testing with Shapiro-Wilk-Test.

